# GW4869 depletes macrophages and increases number of extracellular vesicles in murine peritoneal cavity fluid

**DOI:** 10.64898/2026.01.20.700528

**Authors:** Johann Mar Gudbergsson, Pedro Henrique Gobira, Rodrigo Grassi-Oliveira, Anders Etzerodt

**Author notes:** Correspondence Anders Etzerodt, Ph.D.

## Abstract

Pharmacological approaches to inhibit extracellular vesicle (EV) release in vivo are increasingly used, although the effects of such compounds on local cellular environments are not fully understood. In this study, we examined the impact of intraperitoneal (i.p.) administration of the neutral sphingomyelinase inhibitor GW4869 on EV levels in the peritoneal cavity. Repeated GW4869 i.p. injections elicited a marked local inflammatory response, reduced peritoneal macrophage numbers, and paradoxically increased EV concentrations 24 hours after treatment. Independent macrophage depletion reproduced this rise in EV levels, indicating that macrophage loss and associated cellular remodeling contribute substantially to EV accumulation. These observations indicate that GW4869 can perturb local immune homeostasis in vivo, a confound that must be considered when using this compound as a putative selective inhibitor of EV release.

## Introduction

Extracellular vesicles (EVs) are a collection of heterogeneous lipid bilayer vesicles, including late endosome-derived exosomes and plasma membrane-derived microvesicles. Exosomes are usually described as the smallest EVs, ranging from 30 to 200 nm, whereas microvesicles range from 100 to 1000 nm. Subclasses of EVs are in theory defined by their pathway of biogenesis but often in practice defined by size and/or density, such as large EVs being membrane-derived microvesicles and small EVs being exosomes. The link between EV size and biogenesis is not straightforward, if it exists at all, and consequently we will refer to them as small and large EVs. EVs have been shown to play key roles in health and disease by affecting neighbouring or distant cells with their cargo, and inhibition of EV release has been crucial for understanding their role in cell-to-cell communication [1]. Several small-molecule inhibitors have been shown to affect EV secretion, including calpeptin, manumycin A, imipramine and the neutral sphingomyelinase 2 (nSMase2) inhibitor GW4869 [2]. nSMase2 generates ceramide that is important for intraluminal vesicle formation and exosome secretion, and GW4869 has been instrumental for elucidating role of ceramide in the exosomal pathway [3]. Consequently, GW4869 has become one of the most widely used EV inhibitors and has helped demonstrate how EVs contribute to diverse processes, including viral transmission [4], cell motility [5], education of macrophages [6], and response to chemotherapy [7]. Although GW4869 is commonly described as well tolerated following intraperitoneal (i.p.) administration in mice, systematic assessments of local immune or cellular effects are limited, and reports of its ability to reduce EV release in distant tissues, such as the brain, have relied largely on indirect or heterogeneous measurements [8], [9], [10], [11], [12]. Only two studies have also investigated peritoneal cells or EVs after i.p. injection of GW4869 without mentioning adverse effects on the cellular compartments, indicating that the inhibitor is well-tolerated locally as well [13], [14]. To directly test whether GW4869 reduces EV abundance in vivo, we specifically evaluated its effects on EV levels and cellular composition within the peritoneal cavity. Here, we found that repeated intraperitoneal injections of GW4869 induced local inflammation and selective depletion of resident macrophages. Unexpectedly, rather than reducing EV abundance, we observed a significant increase in peritoneal EV numbers 24 hours after the final injection. Independent macrophage depletion reproduced both the rise in EVs and the influx of inflammatory cells, indicating that changes in the cellular milieu, rather than direct inhibition of EV biogenesis, largely explain this effect. Together, these results demonstrate that GW4869 markedly perturbs local immune homeostasis in vivo, which can secondarily alter EV levels and complicates its use as a selective EV inhibitor.

## Materials and methods

### Animals and housing

Male C57BL/6J mice (8 weeks old; Janvier Labs, France) were group-housed (five per cage) under controlled environmental conditions (22 ± 2 °C; 60 ± 5 % relative humidity) with ad libitum access to food and water. Animals were maintained on a 12 h light/dark cycle (lights on at 06:00 h) and provided with environmental enrichment, including a stainless-steel tunnel shelter, nesting material, and a wooden stick, throughout the study. The in vivo ShredR model was performed using Cas9-EGFP^fl/fl^ (B6J.129(B6N)-Gt(ROSA)26Sor<tm1(CAG-cas9^*^,-EGFP)Fezh>/J) mice bred in-house. These mice were crossed with Csf1r^Cre/+^ (C57BL/6-Tg(Csf1r-cre)1Mnz/J) mice to generate animals with Csf1r-dependent Cas9 expression. All experimental procedures were approved by the Danish National Committee for Ethics in Animal Experimentation (protocol 2023-15-0201-01528 for GW4869 studies and 2021-15-0201-01083 for the ShredR-LNP study) and were conducted in accordance with the European Community Council Directive 2010/63/EU.

### Drug and Treatment Protocol

Prior to the start of drug administration, animals were acclimatized to the housing facility for two weeks. The neutral sphingomyelinase inhibitor GW4869 (TargetMol Chemicals Inc, USA) was prepared as an 8 mg/mL stock solution in dimethyl sulfoxide (DMSO) and freshly diluted in sterile saline to a final DMSO concentration of 2.5 % immediately before use. Mice received daily i.p. injections of GW4869 (2.5 mg/kg) or vehicle (saline + 2.5 % DMSO) for 14 consecutive days, following established protocols [11], [12] . Animals were randomly assigned to one of two treatment groups (vehicle or GW4869) and sacrificed by decapitation either 2 hours or 24 hours after the final injection. Mice were acclimatized for two weeks before receiving daily i.p. (i.p.) injections of GW4869 (2.5 mg/kg) or vehicle for 14 consecutive days. Animals were sacrificed either 2 h or 24 h after the final injection to assess potential temporal effects of nSMase inhibition. Peritoneal lavage samples were collected immediately after sacrifice.

### Macrophage depletion using ShredR lipid nanoparticles

Lipid nanoparticles (LNPs) loaded with ShredR sgRNA were prepared as previously described (see Table 1 for lipid formulation and sgRNA sequences) [15]. Briefly, RNA dissolved in the aqueous phase (100 μM sodium acetate buffer, pH 4.5; Thermo Scientific) was combined with the lipid phase at a flow rate ratio of 4:1 (aqueous to lipid), maintaining a total flow rate of 5□mL/min using the NanoAssemblr^®^ platform. The resulting LNPs were subsequently dialyzed twice overnight against 1xPBS (Cytiva) using 14□kDa MWCO dialysis tubing (Spectra/Por^®^□4). To concentrate the preparation, the dialysis membranes were covered with polyethylene glycol (PEG, Mw□35,000; Sigma-Aldrich) for 10–20□minutes at room temperature. Afterwards, the LNPs were passed through a 0.2□μm cellulose acetate sterile filter (Avantec^®^). Lipid content was quantified by high-performance liquid chromatography (HPLC) using a Dionex Ultimate□3000 system (Thermo Fisher Scientific) equipped with a Nucleosil□100-3C18 column. RNA concentration was assessed using the modified Quant-iT□RiboGreen RNA assay (Thermo Scientific□#R11490) following the Precision□NanoSystems protocol. For storage, LNPs were frozen at□−70□°C in the presence of□5% sucrose (Merck) as a cryoprotectant to ensure RNA stability. For *in vivo* use, LNPs were diluted in sterile PBS and 250 ng LNP (based on RNA concentration) was injected i.p. in 0.2 mL into CD115^Cre^ x Cas9-GFP^fl/fl^ and control mice. Peritoneal cells and fluid were harvested 24 hours later as described below.

**Table 1.**
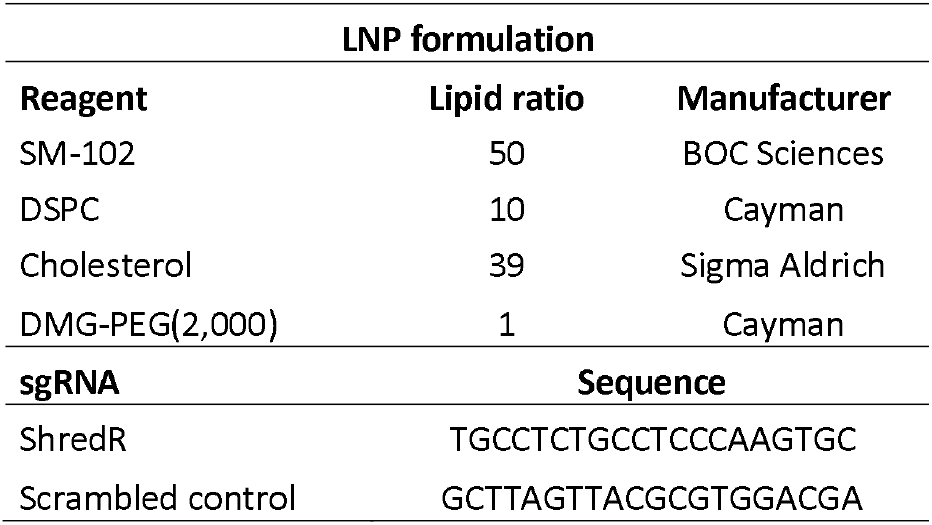
LNP formulation and sgRNA sequences.

### Extraction of peritoneal cells and fluid

After decapitation, mice were injected i.p. with 5 mL of PBS + 2 mM EDTA (Invitrogen, #15575020), and the abdominal region was massaged for 5-10 s prior to extraction of the fluid, as previously described [16]. To extract the fluid, a small incision was made in the skin and blunt dissection done to expose the outer muscle layer. The lateral skin on the right side of the mouse was left attached to allow for stretching of the abdominal muscle layer and displace the abdominal organs with a finger to create an i.p. space for fluid aspiration. A 25G needle was inserted into the peritoneal cavity rostrally pointing caudally and fluid was slowly aspired. Approximately 4.5 mL could be recovered from each mouse. The peritoneal fluid was then centrifuged at 400xg for 5 min at 4C to pellet cells and cells were resuspended in FACS buffer (PBS, 2.5 % FBS, 1 mM EDTA, 0.1 % NaN3) for further analyses. The supernatant was transferred to a new tube and centrifuged for 15 min. at 2,000xg at 4C to pellet cell debris. The resulting supernatant was aliquoted, frozen and stored at -20C until analysis.

### Spectral flow cytometry

Peritoneal cells were counted on a NucleoCounter NC-3000 using A8 NC-slides and viability solution 13 (ChemoMetec). Stain mix was prepared using FACS buffer with mouse Fc-block (clone 2.4G1), brilliant stain buffer (#659611, BD Horizon) and oligo block [17], and antibodies (see antibody panel and dilutions in Table 2). 1 million cells were stained at RT for 30 min, resuspended in FACS buffer + viability dye, and 300-400.000 events were recorded on a SONY ID7000 spectral flow cytometer using standardized mode. Reference spectra were made from antibody single stains using UltraComp eBeads Plus compensation beads (01-3333-42, Invitrogen) and spectra for viability dye was made using ViaComp beads (SSB-07-A, Slingshot Biosciences). Individual spectra were loaded to the sample groups and unmixing was done using the SONY ID7000 Software. After unmixing, sample analyses were done using FlowJo 10.10.0 (BD Life Sciences), and graphs and statistics were done with GraphPad Prism 10 (Dotmatics). Gating strategy for cells can be seen for GW4869 and vehicle treatments in Figure S1.

**Table 2.** Spectral flow panel.

| Primary Excitation laser | Antigen | Clone | Catalogue | Vendor | Fluorochrome | Dilution |
| --- | --- | --- | --- | --- | --- | --- |
| UV 355nm | NK1.1 | PK136 | 564144 | BD Horizon | BUV395 | 200 |
|  | CD102 | 3C4(mIC2/4) | 751392 | BD Optibuild | BUV615 | 400 |
|  | Tim4 | RMT4-54 | 749134 | BD Optibuild | BUV737 | 400 |
|  | CD19 | 1D3 | 568287 | BD Horizon | BUV805 | 400 |
| Violet 405nm | SYTOX Blue |  | S34857 | Invitrogen |  | 10000 |
|  | F4/80 | T45-2342 | 743282 | BD Bioscience | BV650 | 400 |
|  | CD115 | AFS98 | 135515 | BioLegend | BV711 | 400 |
|  | Siglec F | E50-2440 | 740956 | BD Optibuild | BV786 | 400 |
| Blue 488nm | Lyve1 | ALY7 | 53-0443-82 | Invitrogen | AF488 | 800 |
|  | CD45 | 30-F11 | 569257 | BD Horizon | RB545 | 400 |
|  | Ly6C | AL-21 | 571103 | BD Horizon | RB613 | 400 |
|  | CCR2 | 475301 | 757170 | BD Optibuild | RB744 | 400 |
| Yellow 561nm | CD9 | MZ3 | 124805 | BioLegend | PE | 4000 |
|  | CD3e | 145-2C11 | MCA2690C | BioRad | PE-Cy5 | 400 |
|  | CD226 | 10E5 | 567651 | BD Pharmingen | PE-Cy7 | 800 |
| Red 637nm | FOLR2 | 10/FR2 | 153305 | BioLegend | APC | 800 |
|  | Ly6G | 1A8 | 560600 | BD Pharmingen | APC-Cy7 | 200 |
|  | CD11b | M1/70 | 101287 | BioLegend | APC-Fire/810 | 800 |

### Nano flow cytometry

All nano flow cytometry experiments were performed on a CytoFLEX Nano (Beckman Coulter), as previously described [18]. We did not enrich for EVs and kept samples untouched apart from the dilution during extraction of peritoneal fluid. 50 uL of sample was added to a 96-well plate and stained with 50 uL anti-CD9-AF647 antibody (clone MZ3, #124809, BioLegend) diluted to 1 µg/mL in PBS, which was added on top and mixed. Prior to making antibody stain mix, all antibody vials were centrifuged at 16,000xg for 5 min to avoid/reduce antibody aggregates. Plate was incubated overnight at 4C with antibody. The next day, samples were stained at 1X concentration with a mix of the membrane dyes ExoBrite 410/450 CTB (#30111-T, Biotium) and ExoBrite 515/540 True EV membrane stain (#30129-T, Biotium) which was diluted in PBS and 50 uL added to the samples and incubated 1-2 hours at RT. Just before analysis samples were diluted in PBS to a final dilution of 1:200 and run on the CytoFLEX Nano. A vehicle (PBS) control sample was run to set the vSSC1 threshold which in our setup was set at 300 in all experiments. Scatter detectors gains were kept according to QC values, but relevant fluorescent detectors were maxed to 3000. Single stain and unstained controls were run to setup compensation. PBS + stain control was run as a background control and for all quantifications the background values were subtracted using the PBS + stain control. A detergent control was used where 2 % Triton X-100 (CAT) was prepared and sterile filtered, and added 1:1 to an already analysed sample and incubated for 30 min at RT. The sample was then analysed to show that the particles that fall into our EV gate are in fact sensitive to detergents. Scatter calibration was done as previously described [18], [19], [20]. We have compiled representative controls in Figure S2. Compensation and data analysis was done in CytExpert Nano (Beckman Coulter) and FlowJo™ v10.10.0 (BD Life Sciences) and quantitative graphs and statistical analyses were created in GraphPad Prism 10 (Dotmatics).

## Results

### GW4869 EV inhibitor depletes macrophages and causes inflammation in the mouse peritoneal cavity

In an attempt to inhibit EV release from mouse peritoneal cells, we treated mice i.p. for 14 consecutive days with the nSMase2 inhibitor GW4869 or vehicle (Figure 1A). To analyse the effect on EV release and cellular dynamics we harvested peritoneal cells and EVs 2 and 24 hours (h) after the last injection. Surprisingly, we observed a specific reduction of total cells in GW4869 treated mice 24 h after last injection compared to vehicle control. In contrast, there was no major difference in cell viability, nor did we observe any difference in cell numbers 2 h after last (Figure 1B, C). This observation warranted a deeper phenotypic analysis of the peritoneal cells after GW4869 treatment using spectral flow cytometry to detect any changes in cell populations. Strikingly, we observed that GW4869 treatment specifically depleted peritoneal cavity of macrophages at both timepoints (Figure 1D-E, Figure S3A). Further analysis showed that both large peritoneal macrophages (LPM) and small peritoneal macrophages (SPM) where depleted after treatment, as remaining F4/80+CD11b+ macrophages did not express canonical LPM markers Icam2/CD102 or the SPM (CD226) markers (Figure 1F-G). Simultaneously, a high neutrophil and eosinophil influx was seen 2 hours after treatment, along with a significant increase in CD11b MFI indicating an inflammatory environment (Figure 1E, H). The number of neutrophils and eosinophils and their respective CD11b MFI was reduced after 24 hours, indicating a strong and short-lived response to GW4869 (Figure 1E, H). In summary, we found that the nSMase2 inhibitor GW4869 eliminates the macrophage compartment in the peritoneal fluid and creates an inflammatory environment with activated neutrophils and eosinophils.

**Figure 1.**
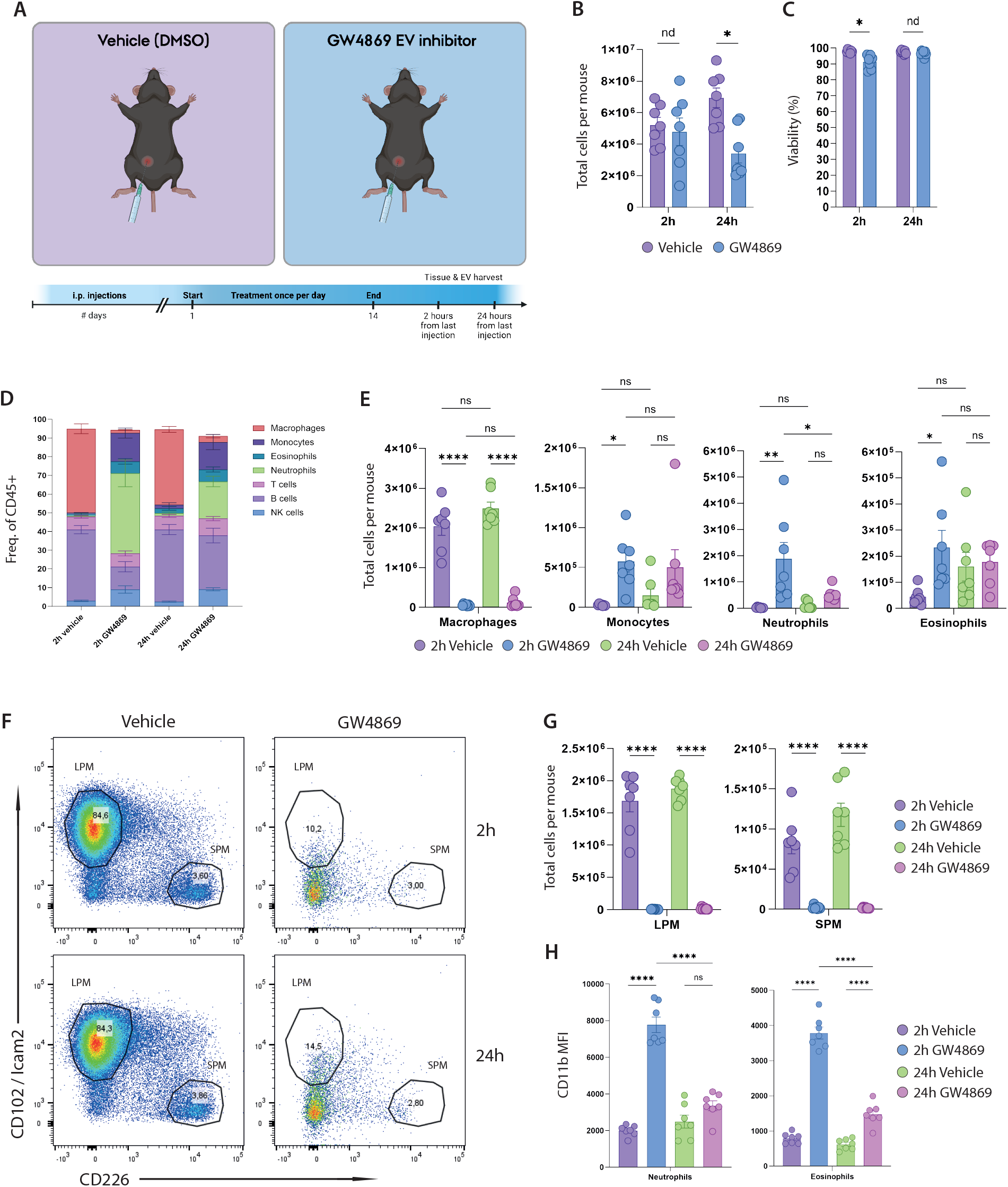
GW4869 depletes peritoneal macrophages and induces local inflammation. A) Schematic overview of treatment schedule. B) Total peritoneal cells per mouse (p = 0.00151). C) Viability of cells (p = 0.00154). D) Stacked bar charts of cell types represented as frequency of CD45^+^ cells. E) Bar charts of total number of macrophages (p = <0.0001), monocytes (p = 0.0385), neutrophils (p = 0.0017, p = 0.0282) and eosinophils (p = 0.036). F) Representative bivariate plots of LPM and SPM populations. G) Bar chart showing total LPMs (p = <0.0001) and SPMs (p = <0.0001) per mouse. H) CD11b MFI of neutrophils (p = <0.0001) and eosinophils (p = <0.0001). n = 7.

### GW4869 increases number extracellular vesicles in mouse peritoneal fluid

After assessing the consequence of GW4869 treatment on the cellular compartment, we examined the effects of the nSMase2 inhibitor on particles and EV release in the peritoneal fluid at 2 h and 24 h after last treatment by nano-flow cytometry using the CytoFLEX Nano from Beckman Coulter (Figure 2A). All samples were analyzed on the same day, and an inclusion gate was made based on the dilution buffer (freshly opened, sterile PBS). We quantified all particles within the inclusion gate and found a drastic increase in the GW4869 treated mice 2 h after treatment, whereas the particle numbers were normalized again at 24h (Figure 2B, C). To specifically look at single EVs, the samples were stained with a lipophilic dye (ExoBrite True EV Membrane stain) and a cholera toxin b conjugate (ExoBrite CTB). Surprisingly, we observed that the total number of EVs was similar between treatment groups 2 h after treatment but was increased ∼2-fold after 24 h in the GW4869 treated mice (Figure 2D). This effect was reflected in CD9^+^ EVs (Figure 2E), whereas the total number of CD9^+^ cells in the peritoneal cavity was significantly decreased at 24 h after treatment (Figure 2F). However, as shown in Figure 1, an influx of inflammatory eosinophils and neutrophils occurs due to GW4869 treatment, and they are all CD9^+^ (Figure S3B). Furthermore, eosinophils and neutrophils from treated animals also had increased CD9 expression compared to vehicle (Figure S3C). We observed an increase in median size of particles in the EV gate from treated animals at both 2 h (145.8 nm vs 175.9 nm) and 24 h (142 nm vs 165 nm), but not for particles in the inclusion gate or CD9+ EV gate (Figure 2G). Lastly, we repeated the experiment at 24 h and got similar results (Figure S4). In summary, we find that the EV inhibitory effects of GW4869 seem to be inefficient in the mouse peritoneal cavity at the dose used here, and instead highly increases the presence of particles and EVs, likely as a consequence of macrophage depletion and influx of other activated immune cells.

**Figure 2.**
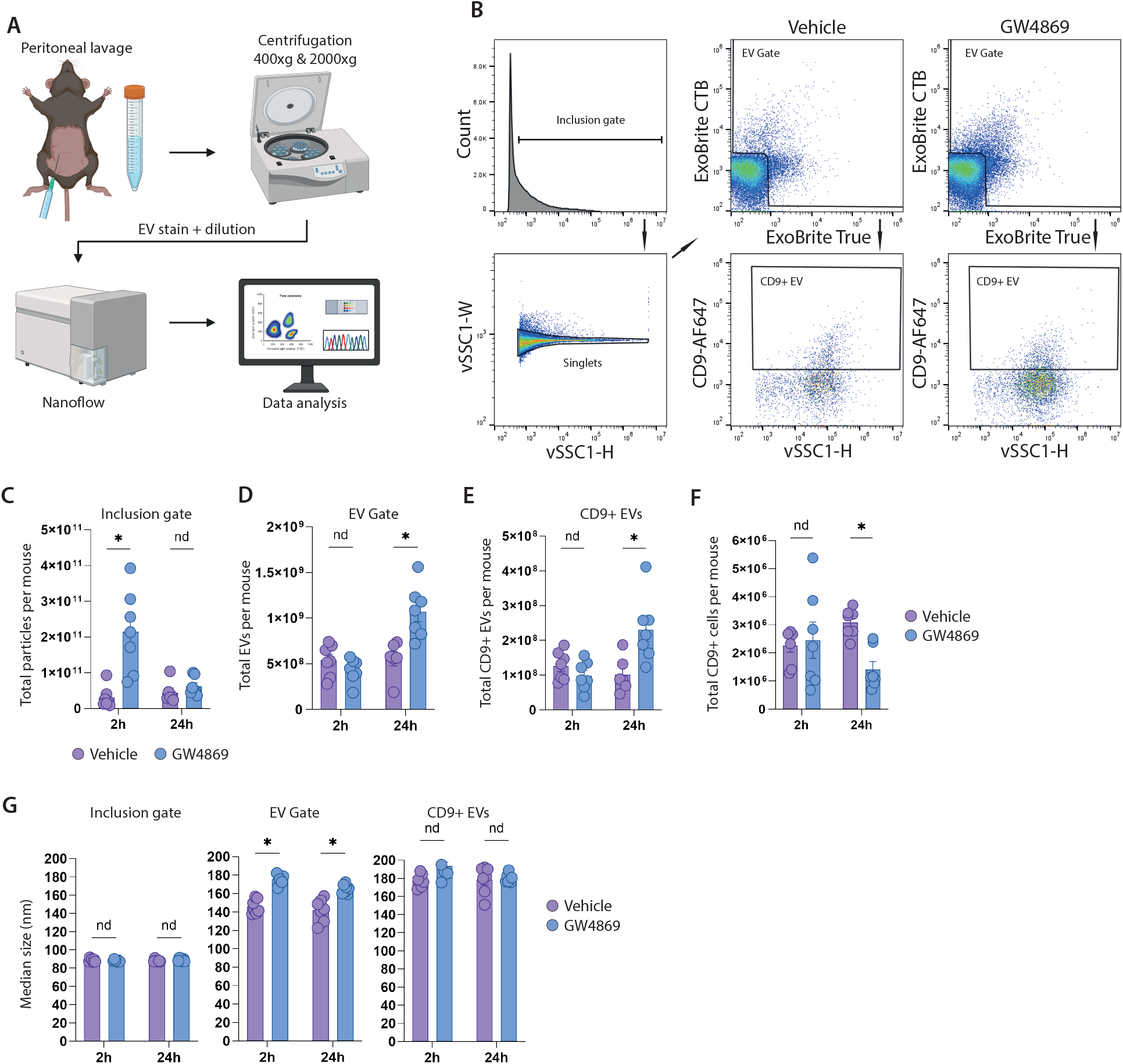
GW4869 increases number of total peritoneal EVs. A) Brief schematic overview of sample workflow. B) Gating strategy for nano flow cytometry analyses peritoneal particles and EVs. C) Bar chart of total particles per mouse derived from inclusion gate (p = 0.00142). D) Total EVs per mouse (p = 0.00338). E) Total CD9^+^ EVs per mouse (p = 0.01146). F) Total CD9^+^ cells per mouse (p = 0.00026). G) Median size in nm derived from size calibration using FCMPASS within inclusion gate, EV gate (p = <0.0001, p = 0.00062), and CD9^+^ EVs. n = 7.

### Depletion of peritoneal macrophages increases the number of extracellular vesicles

Since GW4869 treatment resulted in loss of peritoneal macrophages and increase in EVs numbers, we next wanted to test if similar effect would be seen when specifically depleting peritoneal macrophages using a targeted approach. To this end we bred Csf1r-Cre mice with Cas9-egfp floxed mice to generate Cas9-expressing macrophages in the peritoneal environment and delivered SM102 lipid nanoparticles containing our cell-depleting ShredR sgRNAs (Figure 3A), as previously described [15]. Although ShredR treatment led to an increase in total peritoneal cells with no difference in cell viability (Figure 3B-C), flow cytometric analysis confirmed an efficient depletion of macrophages after 24h. As hypothesized, the depletion of LPMs and SPMs was accompanied by an influx of monocyte and neutrophil as previously observed using GW4869 (Figure 3D-F). Interestingly, although total number of particles present in the peritoneal fluid was not different between the treatments, ShredR treatment led to a specific increase in total EVs and total CD9^+^ EVs similar to what was observed for GW4869 treatment (Figure 3G-I). These results indicate that a depletion of macrophages from the peritoneal cavity fluid results in an inflammatory environment which is accompanied by an increased number of EVs.

**Figure 3.**
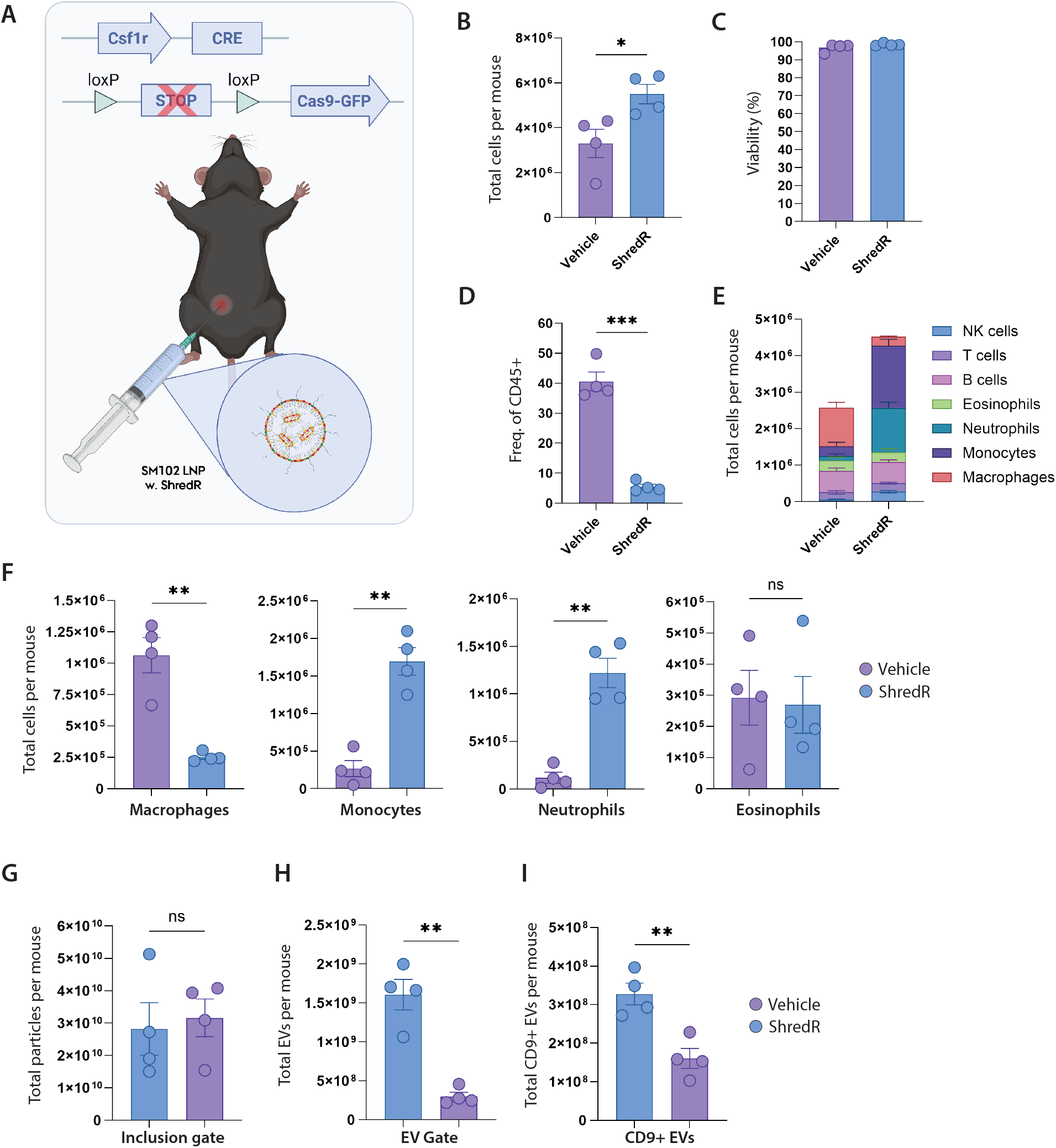
Macrophage depletion increases number of total peritoneal EVs. A) Illustration of mouse model and treatment applied. B) Total peritoneal cells per mouse (p = 0.0332). C) Viability of cells. D) Bar chart showing macrophage frequency of CD45^+^ cells (p = 0.0009). E) Stacked bar charts of cell types represented as frequency of CD45^+^ cells and as total cells per mouse. F) Bar charts representing total macrophages (p = 0.0096), monocytes (p = 0.0013), neutrophils (p = 0.0031), and eosinophils per mouse. G) Bar chart of total particles per mouse derived from inclusion gate. H) Total EVs per mouse (p = 0.005). I) Total CD9^+^ EVs per mouse (p = 0.0048). n = 4.

## Discussion

The effects of GW4869 on immune cell subsets in vivo has not been extensively characterized. In our study, examining its impact on mouse peritoneal cells, we observed an almost complete depletion of peritoneal macrophages. This depletion resulted in local inflammation with increased neutrophil and eosinophil influx, and instead of reduction of EV abundance, we saw a two-fold increase in total EVs 24h after the final treatment. The inclusion of two time points (2 h and 24 h) allowed for the assessment of potential temporal effects of nSMase inhibition, controlling for confounding influences related to the prolonged 24 h drug-free interval and potential peak enzymatic activity of neutral sphingomyelinase. At the 2 h time point we saw no difference in total EVs but a vast difference in total particle numbers. An increase in EVs after GW4869 treatment has not, to the best of our knowledge, been reported before, but previous studies showed that GW4869 treatment specifically reduced exosome release and increased MV/apoptotic vesicle release [21], [22]. This conclusion is only based on increased number of particles (analyzed by NTA) that precipitate at medium-speed centrifugation (10-14,000xg) after GW4869 treatment compared to high-speed (>100,000xg). However, such analysis cannot properly distinguish between exosomes and microvesicles due to their overlap in size and density [2], [21], [23]. Interestingly, one of the studies showing inhibitory effect of GW4869 on EV release also reported an increase in apoptotic-related Annexin A5 expression in the 10,000xg fraction, indicating a decrease in cellular health after GW4869 treatment [22].

We could only find one study that investigated the peritoneal microenvironment after GW4869 treatment, and they did not report any adverse effects on the peritoneal cells but did see a decrease in EVs at 2.5 mg/kg dosage, similar to the dose we used [13]. It is important to note that this study only gave a single injection of GW4869 compared to our 14-day treatment schedule, which might be gentler on the macrophages. Another important point is that they used a Zymosan A-induced sterile peritonitis model, which has previously been shown to deplete macrophages from the peritoneal fluid and cause a general increase in sphingolipid synthesis [24], [25], [26]. Whether pre-depletion of large peritoneal macrophages by sterile peritonitis before GW4869 injection results in fewer effects on cellular function is not known, but this would be an interesting topic to investigate in the future.

The impact of GW4869 on EV numbers has largely been ascribed to reduced EV release, rather than to potential cellular consequences such as depletion of specific cell populations or alterations in cellular lipid composition [27], [28], [29]. Nevertheless, inhibition of nSMase2 has been reported to perturb fundamental cellular processes, including TNF signaling and autophagy-related pathways [30], [31]. Because ceramide is a key mediator of inflammatory signaling and has been implicated in inflammasome assembly and IL-1β secretion, pharmacological nSMase2 inhibition is likely to influence immune cells such as macrophages [32], [33], [34]. Consistent with this, GW4869 has been shown to exert anti-inflammatory effects in several disease models, with evidence for direct actions on macrophages [35], [36], [37]. Inflammation has also been reported to enhance EV release from cultured cells and to associate with higher circulating EV levels in adults with obesity-related adipose tissue inflammation [38], [39], [40]. In this context, the observation that GW4869 depleted macrophages from the peritoneal cavity and induced inflammation raises the possibility that it could simultaneously attenuate inflammatory EV production by infiltrating cells. EV abundance was unchanged 2 h after the final treatment but increased approximately 2-fold at 24 h, a time-dependent pattern that may be compatible with waning GW4869 activity. In parallel, the recognized contribution of macrophages to EV clearance and turnover suggests that loss of these cells could further influence EV accumulation in the peritoneal fluid [41], [42]. Together, these observations underscore the need for further studies on how pharmacological EV inhibitors reshape local cellular environments and, in turn, influence measured EV abundance.

### Concluding remarks

In conclusion, repeated intraperitoneal administration of GW4869 was associated with local inflammation, selective loss of resident peritoneal macrophages, and an unexpected increase in EV abundance 24 h after the final injection. Similar changes were observed after independent macrophage depletion experiments which support the notion that alterations in cellular composition and inflammatory status of the peritoneal cavity, rather than direct inhibition of EV biogenesis alone, might contribute to the elevated EV levels. Together, these findings indicate that GW4869 modulates EV dynamics in vivo through combined effects on EV production and the immune microenvironment, underscoring that secondary immunological changes and tissue context should be carefully considered when employing GW4869 as a pharmacological EV inhibitor and when interpreting in vivo EV measurements.

## Acknowledgement

The authors wish to acknowledge the FACS Core Facility and Laboratory Animal Core Facility at Aarhus University. The CytoFLEX Nano was a generous gift from the Carlsberg Foundation (Grant CF23-1020). The 5-laser ID7000 was a generous gift from the Novo Nordisk Foundation, grant number NNF210C0066798. A.E. was funded by Aarhus University Research Fund, the Novo Nordisk Foundation (NNF20OC0065510 and NNF22OC0080192) and the Danish Cancer Society (R302-A17596). R.G.O was funded by the Independent Research Fund Denmark (DFF, grant no. 3166-00139B). We thank Martin K. Thomsen and Felicity M. Davis (Aarhus University) for generously providing the Cas9-EGFP^fl/fl^ and Csf1r^Cre^ mouse lines.

## FIGURE LEGENDS

**Figure S1.**
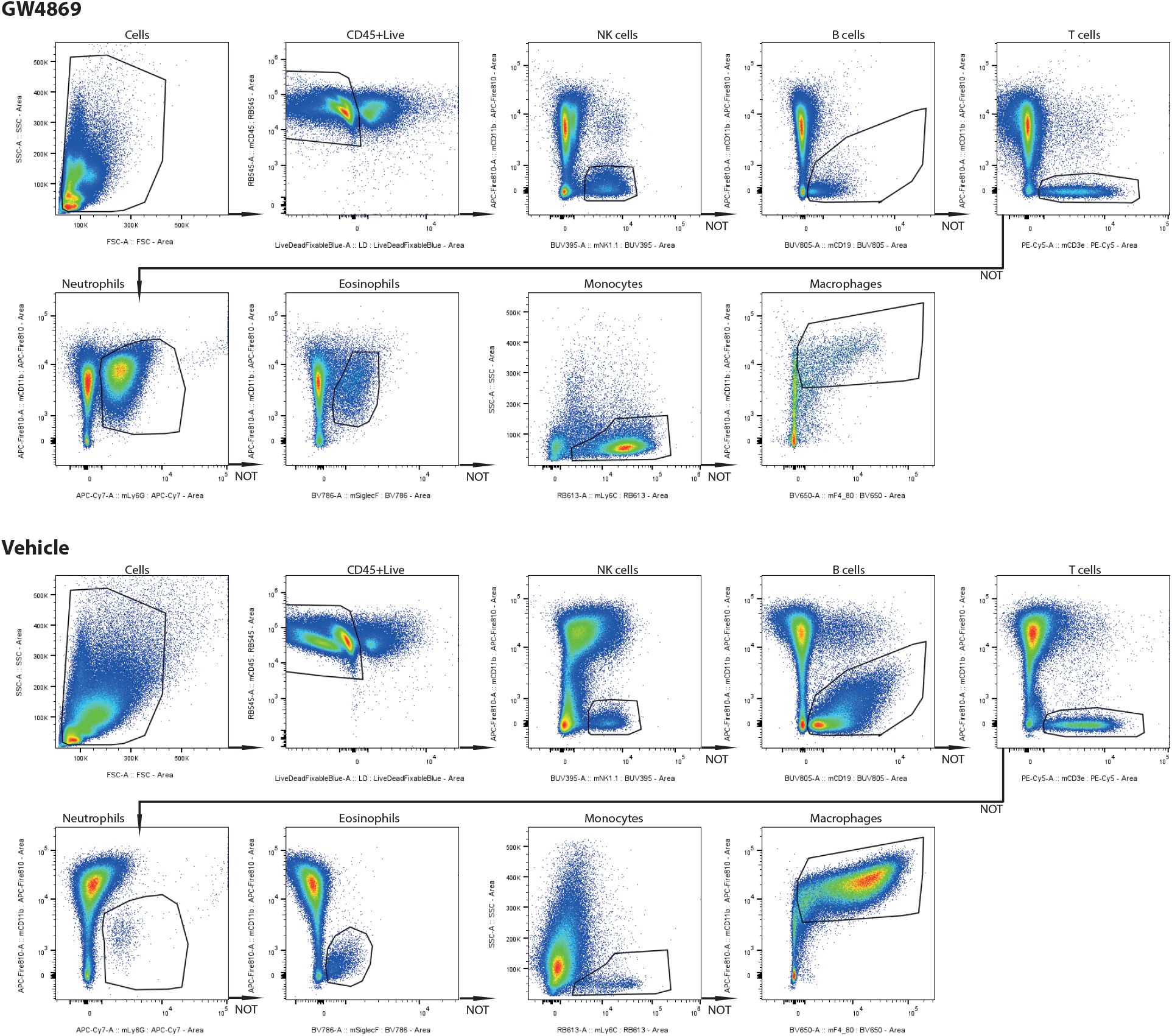
Gating strategy used for spectral flow analysis of cells. Plots are representative for each treatment group.

**Figure S2.**
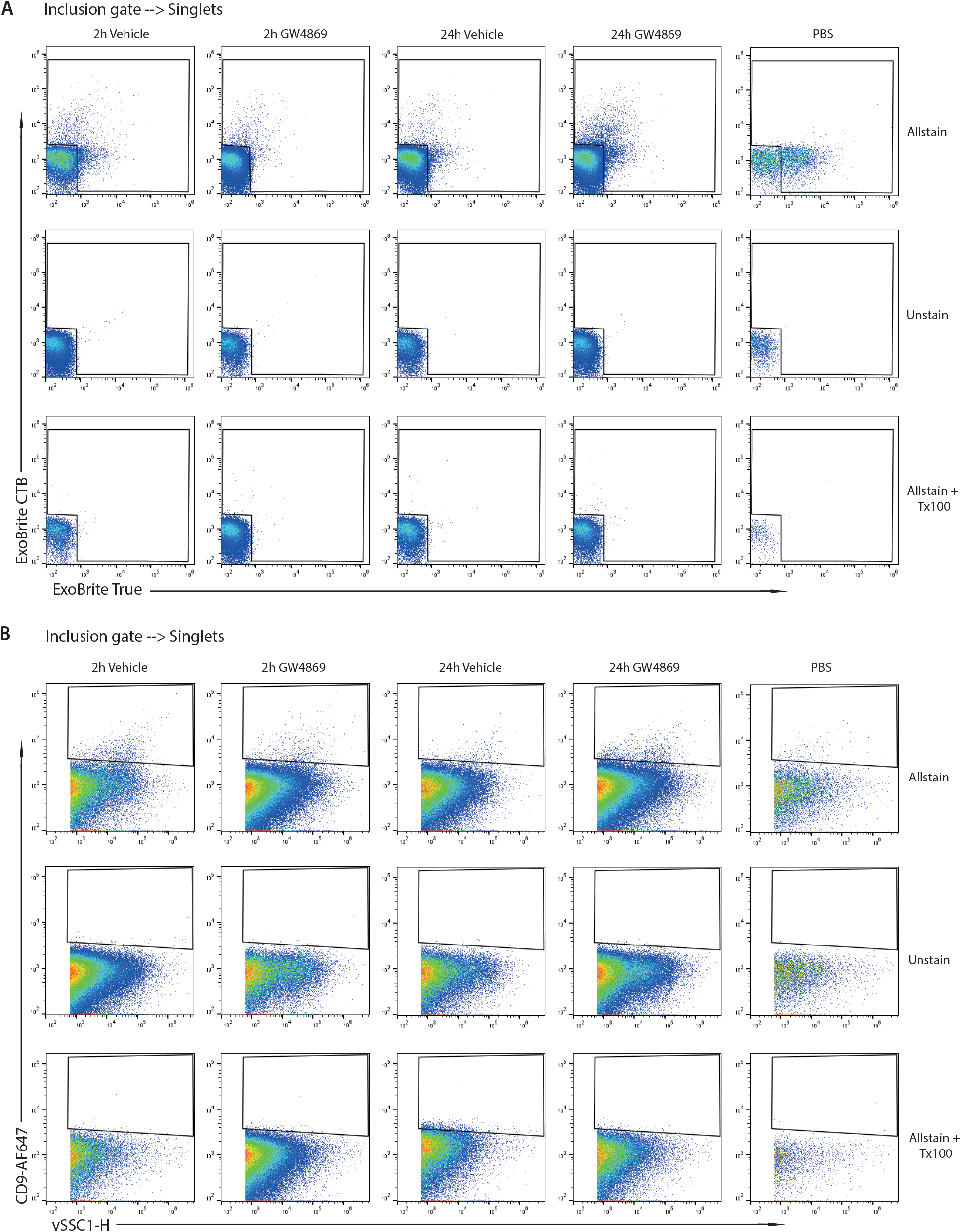
Nano flow cytometry controls. A) Representative controls for EV staining using ExoBrite CTB and ExoBrite True EV Membrane stain. B) Representative controls for CD9-AF647 staining.

**Figure S3.**
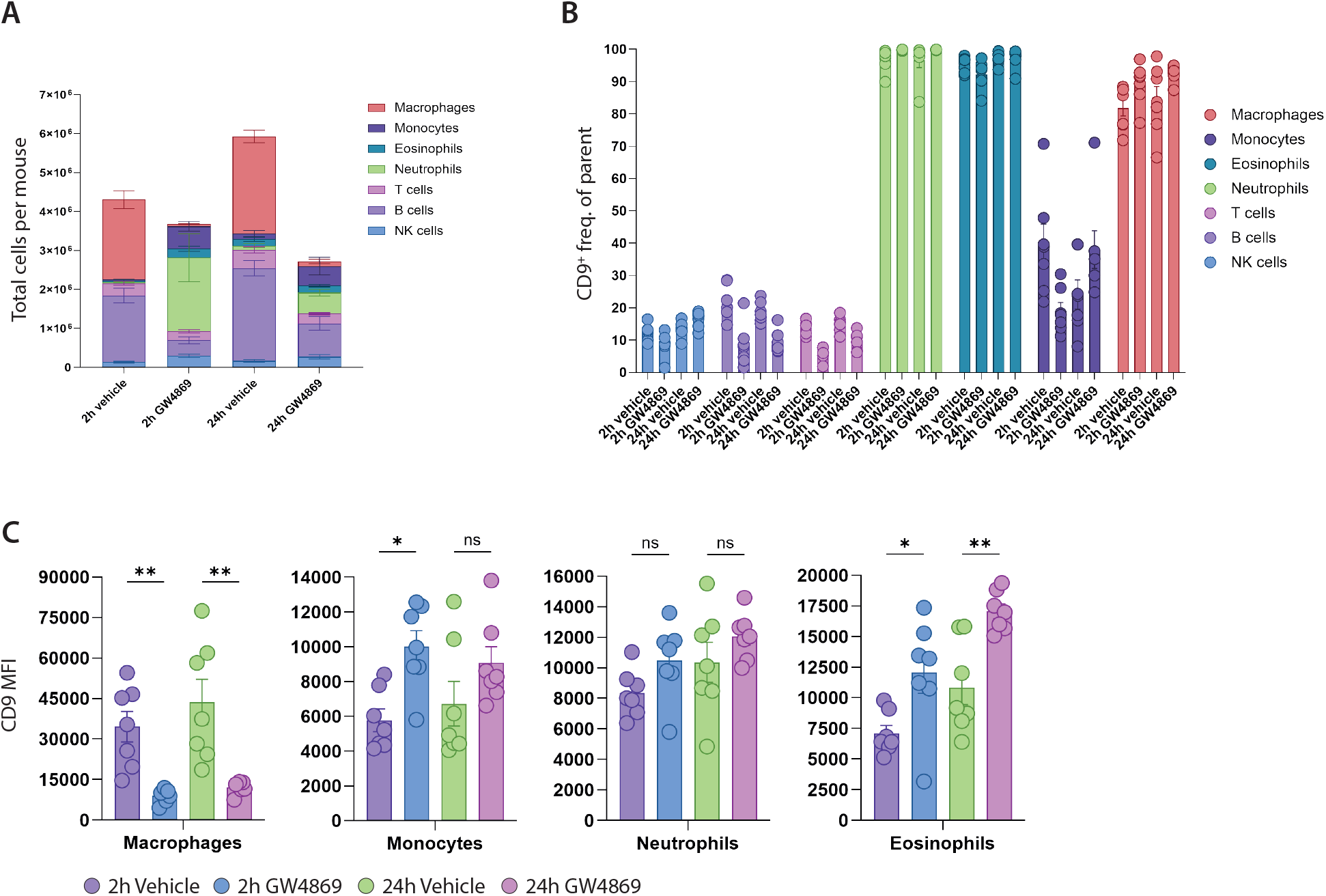
A) Stacked bar charts of cell types represented as total cells per mouse. B) Percentage of CD9^+^ cells within each defined cell type. C) Bar charts displaying CD9 MFI for macrophages (p = 0.0084, p = 0.0011), monocytes (p = 0.0242), neutrophils, and eosinophils (p = 0.0344, p = 0.0063).

**Figure S4.**
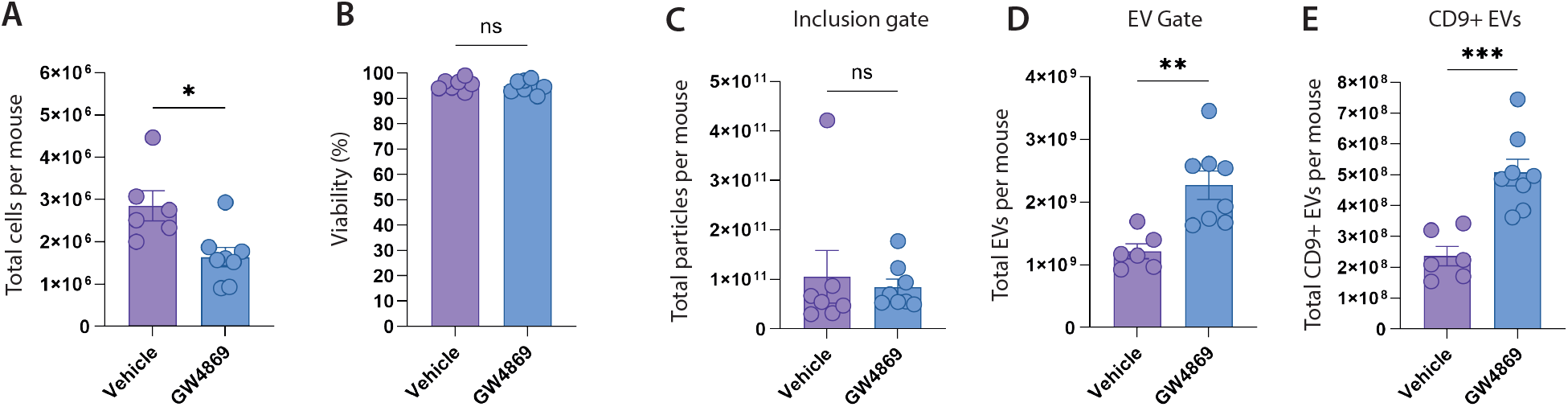
Repeat experiment of GW4869 i.p. 24 h time point. A) Total peritoneal cells per mouse (p = 0.0102). B) Viability of cells. C) Bar chart of total particles per mouse derived from inclusion gate. D) Total EVs per mouse (p = 0.0028). E) Total CD9^+^ EVs per mouse (p = 0.0005). n = 8.

